# MOLECULAR CHARACTERIZATION AND PROMOTER ELEMENT ANALYSIS OF GROWTH REGULATOR BRAGRF5 IN *BRASSICA CAMPESTRIS* L. SSP. *CHINENSIS*

**DOI:** 10.1101/2024.10.13.618120

**Authors:** Xiu-Ying Wang, Jing Zhang, Xian-Hui Qi, Gai-Zhen Li, Chuang-Ye Lan, Qi-Zhang Li

## Abstract

The transcriptional regulation of eukaryotic gene expression is a critical aspect, with transcription factors playing a key role by combining with cis-acting elements in the promoter regions. Growth regulating factors (GRFs) are plant-specific transcriptional factors. The *Arabidopsis thaliana* growth-regulating factor gene (*AtGRF5*), when ectopically expressed in crops such as *Beta vulgaris* (sugar beet) and *Brassica napus* (canola), has an effect of accelerating shoot organogenesis and enhancing the efficiency of genetic transformation. However, an efficient plant regeneration and genetic transformation system in *Brassica campestris* L. ssp. *chinensis*, a major vegetable crop in China, remains to be established. In this work, we identified and characterized the *AtGRF5* ortholog in *B. campestris*, named as *BraGRF5*, from BRAD database. Molecular analysis of *BraGRF5* gene and exploration of its protein interaction network were further conducted. Therefore, the promoter region of *BraGRF5* was thoroughly examined to determine cis-acting elements of various kinds and quantities. It is expected to provide a foundation for regulation of *BraGRF5*, and a informative reference for the improvement of *B. campestris* transformation efficiency and molecular breeding.

## Introduction

Chinese cabbage (*Brassica campestris L*. ssp. *pekinensis*) is one of the most economically significant vegetable in China, renowned for its diverse varieties and substantial market demand and serves as a staple in winter cuisine, and is extensively cultivated throughout the country, particularly in northern regions. Furthermore, morphological classifications include heading and non-heading types, with the former encompassing varieties such as Shandong and Tianjin cabbage, while the latter includes types like bok choy (Liu *et al*., 2018). This rich diversity not only satisfies diverse consumer preferences but also contributes to the sustainable development of the Chinese cabbage industry.

In the regulation of growth and development in Chinese cabbage, growth-regulating factors (GRF proteins) play a pivotal role. GRF proteins are a class of essential transcription factors that influence various aspects of plant growth, including cell expansion, division, and tissue differentiation, which was initially studied in *Oryza sativa* (van der Knaap *et al*., 2000), and subsequently, also identified in *A. thaliana* (Kim *et al*., 2003), *Triticum aestivum* (Huang *et al*., 2021), *Glycine max* (Chen *et al*., 2019), *etc*. Recent studies have highlighted the dual function of GRF proteins in promoting plant growth and enhancing stress resistance, making them crucial for the molecular breeding of commercially valuable crops like Chinese cabbage. In *A. thaliana*, the *GRF4* (*AtGRF4*) has been found to control leaf cell proliferation and cotyledon development (Kim & Lee, 2006), while in *Solanum lycopersicum, SlGRFs* exhibit preferential expression in meristematic tissues and flower buds (Khatun *et al*., 2017).

Significant advancements have been made in elucidating the role of GRF proteins, with researchers utilizing genetic engineering techniques to modify the growth characteristics of Chinese cabbage, resulting in improved yield and quality thereby further enhancing its economic potential (Ou *et al*., 2022). The application of GRF proteins in molecular breeding offers innovative strategies for the enhancement of Chinese cabbage varieties. Through the manipulation of GRF gene expression, researchers can develop high-quality cultivars that are optimally adapted to a wide range of environmental conditions (Zhang *et al*., 2023). These improved varieties not only demonstrate enhanced disease resistance and stress tolerance but also cater to market demands for smaller, fresher products, thereby bolstering their competitive advantage. As consumer preferences for high-quality Chinese cabbage continue to grow, the significance of GRF proteins in breeding programs becomes increasingly pronounced, providing a robust scientific foundation for the sustainable advancement of the Chinese cabbage industry. *B. campestris*, a widely cultivated vegetable crop in China, encounters substantial challenges due to the lack of an efficient plant regeneration and transgenic system (Li *et al*., 2017). In this study, we aim to identify the *AtGRF5* ortholog in *B. campestris*, analyze its fundamental molecular characteristics and protein interaction networks, and investigate the types and quantities of cis-acting elements present in its promoter regions. These efforts will lay the groundwork for understanding and regulating the expression of this target gene. This comprehensive approach is expected to contribute valuable insights for molecular genetic research and facilitate the development of improved breeding strategies for *B. campestris*.

## Material and Methods

### Acquisition and analysis of *B. campestris* GRFs data

The Gene ID and protein sequence of *GRF* family in *B. campestris* were sourced from the Brassicaceae Database (BRAD) (http://brassicadb.cn/); whereas the *AtGRF5* protein sequence of *A. thaliana* was obtained from the NCBI database. The sequence information was then compiled into a fasta format document. In order to identify the homologous sequence *AtGRF5* in *B. campestris*, several sequence alignments were carried out using the NCBI COBALT to build a phylogenetic tree.

### Basic characteristics analysis

Utilizing the Expasy ProtParam instrument (http://web.expasy.org/protparam), the physicochemical characteristics of BraGRF5 were examined. The transmembrane structure of BraGRF5 was analyzed with TMHMM Server v.2.0 (https://services.healthtech.dtu.dk/service.php?DeepTMHMM). The protein secondary structure of BraGRF5 was identified with SOPMA (https://npsa-prabi.ibcp.fr/cgi-bin/secpred_sopm.pl). The conserved domains were conducted by NCBI online tool Batch CD-Search. The Predict Protein (https://predictprotein.org/) was used for protein subcellular localization prediction.

### Expression profiles analysis, Gene Ontology (GO) annotations, and interaction network prediction

The gene expression profiles of *BraGRF5* in different tissues and GO annotations were sourced from BRAD. The STRING was employed to predict the proteins interact with BraGRF5 and further to construct the protein-protein interaction network.

### Chromosome localization and genome structure construction

The BRAD JBrowse (http://brassicadb.cn/#/SearchJBrowse/) was employed for chromosome localization and genome structure construction.

### Promoter sequence analysis

The BRAD Genomic sequence retrieval was used to extract a 2,000 bp fragment upstream of the *Bra027384* start codon as the promoter sequence. Subsequently, the Neural Network Promoter Prediction online software of Berkeley Drosophila Genome Project (https://www.fruitfly.org/seq_tools/promoter.html) was utilized to predict active regions within the promoter. PlantCARE (http://bioinformatics.psb.ugent.be/webtools/plantcare/html/search_CARE.html) was used to analyze cis-acting elements. (Lescot *et al*., 2002).

## Results

### Ortholog identification of BraGRFs and AtGRF5

17 BraGRFs were identified from BRAD, and then employed to aligned with *AtGRF5* (GenBank Accession: AEE75445), using NCBI COBALT tool. subsequently, the evolutionary tree was constructed with default parameters. Results revealed that Bra021521 and Bra027384 clustered together in the same phylogenetic clade with AtGRF5 (Figure 1), showing the highest sequence identity.

**Fig. 1.**
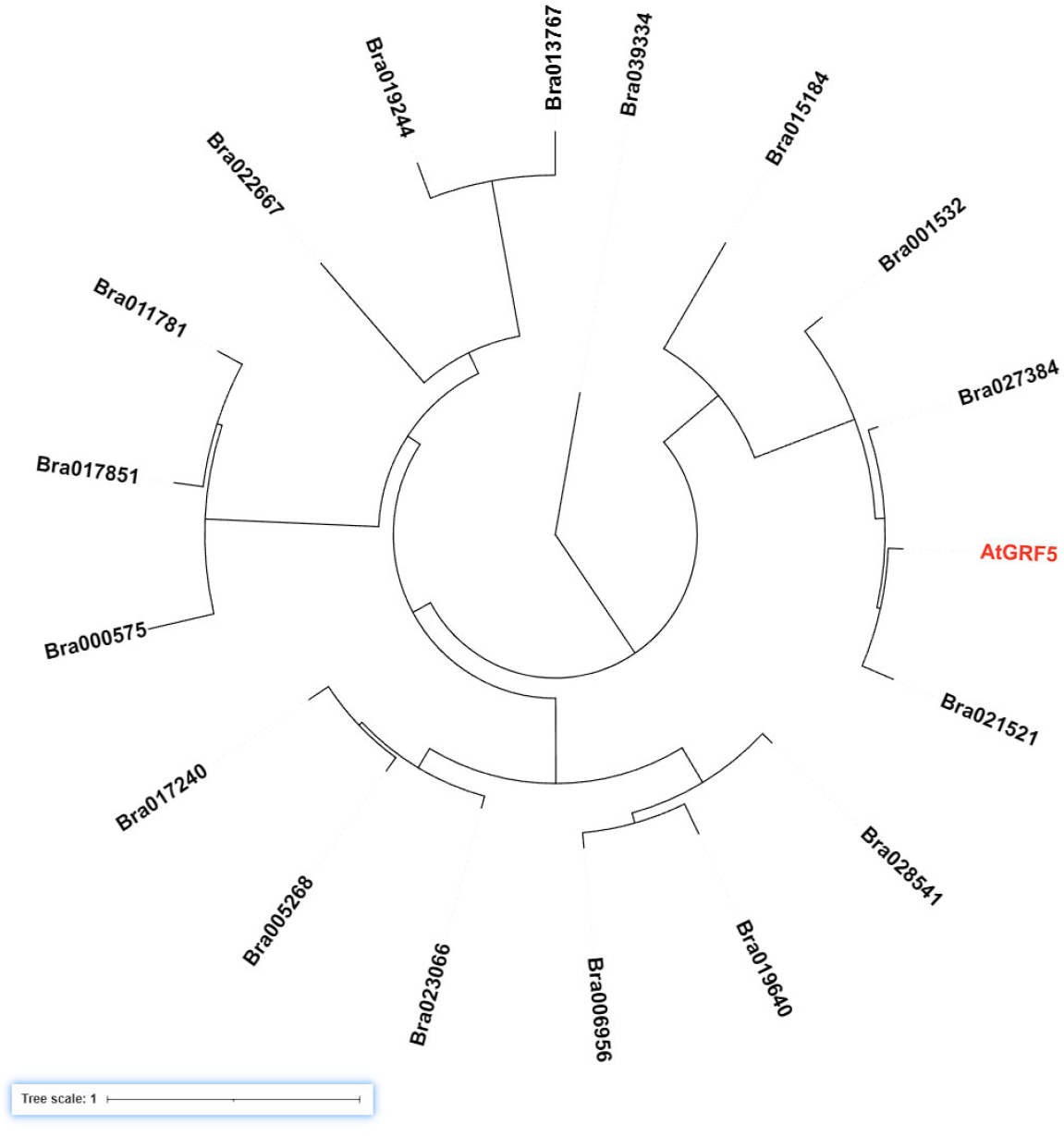
Phylogenetic tree of BraGRFs and AtGRF5 amino acid sequences. The phylogenetic tree was constructed based on full-length aa sequences of 17 BraGRFs proteins with AtGRF5 using NCBI COBALT. The AtGRF5 is marked in red.

Multiple sequence alignment of Bra021521 and Bra027384 with AtGRF5 revealed high amino acid sequence identities between Bra027384 and AtGRF5, while there was a substantial difference at N- and C-terminal of Bra021521 and AtGRF5 (Figure 2). To screen BraGRF5 more accurately, subsequent examination of the conserved structural domains was conducted, results were consistent with that, in agreement with GRF5 in several species, including *Ipomoea Batatas, Cocos nucifera, Zea mays, Dendrobium catenatum*. In particular, similar to AtGRF5 in *A. thaliana*, the C-terminal region of Bra027384 contains the conserved GRF structural domains QLQ and WRC, located at amino acids 16-50 and 82-121, respectively; whereas Bra021521 only contained WRC domain and lacked QLQ domain. The conserved Gln-Leu-Gln residues in the QLQ domain, containing QX_3_LX_2_Q motif, plays a crucial role in mediating functional interactions between proteins. QLQ interacted with GRF-Interacting factor 1 (GIF1) to form potent transcriptional activators, playing a role in plant growth and development. The plant-specific WRC domain, known for its conserved Trp-Arg-Cys pattern, contains a zinc-finger motif for DNA binding and a functional nuclear localization signal (Liu *et al*., 2015). Based on this, Bra027384 and AtGRF5 share similar functions, while the function of Bra021521 appears relatively simple. Therefore, Bra027384 has been designated as BraGRF5.

**Fig. 2.**
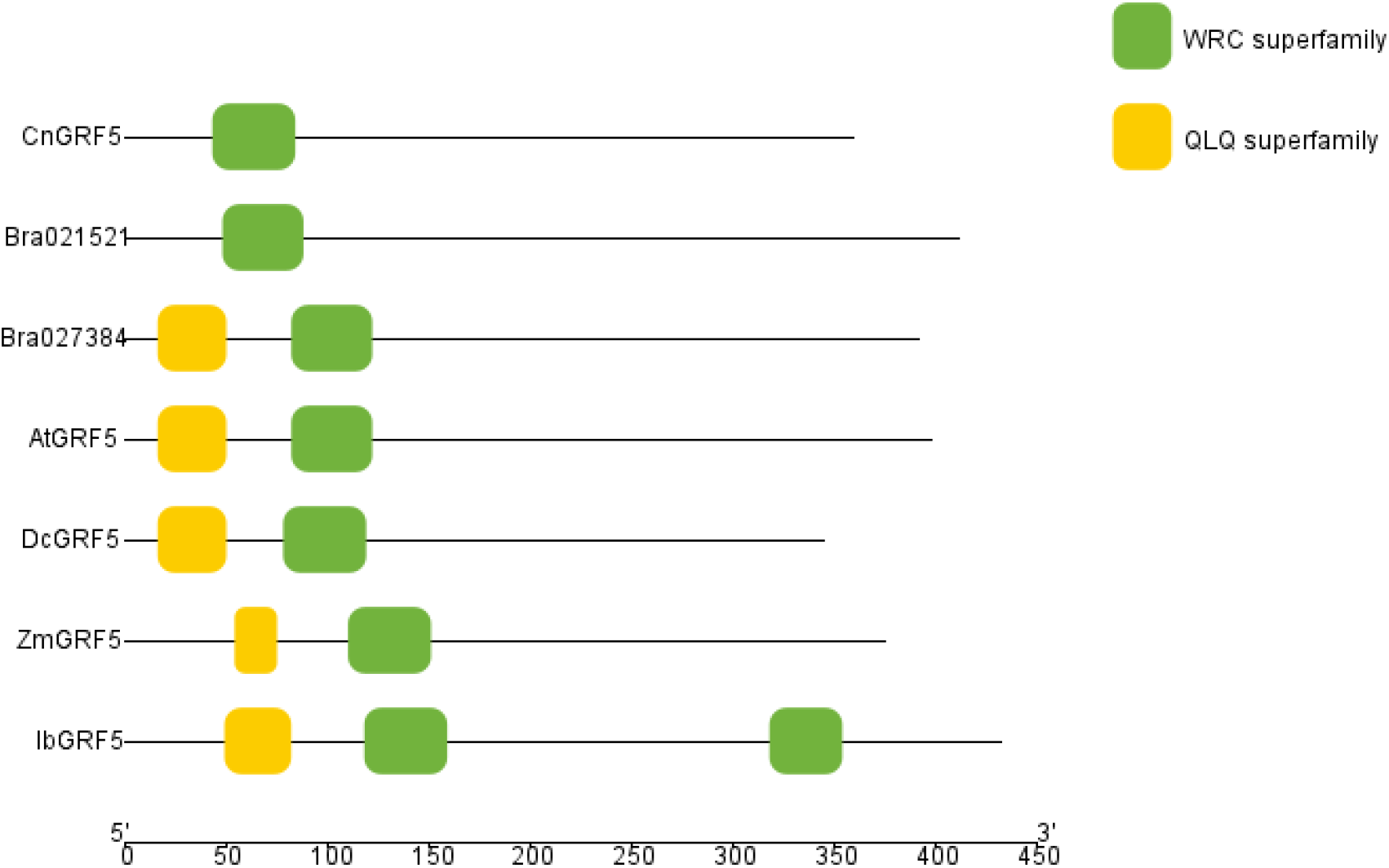
Domain analysis of BrGRFs with other GRF5 proteins from various species. The yellow and green box indicates the conserved QLQ and WRC domains, respectively.

### Molecular characterization of BraGRF5

The protein sequence of BraGRF5 (Bra027384) comprised of 391 amino acids, with a molecular weight of 44,082.15 g/mol and a molecular formula of C_1898_H_2890_N_584_O_611_S_13_. With an instability coefficient of 60.66, it is classified as an unstable protein. The measured abundance of negatively charged residue (Asp + Glu) was 38, and that of positive charged residue (Arg + Lys) was 40. The grand average of hydropathicity (GRAVY) was -1.019, suggesting this protein was hydrophilic. Its aliphatic index was 46.68, and the theoretical isoelectric point (pI) was 8.23.

### Analysis of subcellular localization and secondary structure

BraGRF5 was predicted to localize to the nucleus and did not possess any transmembrane structure. The 84 amino acids (21.48%) formed α-helices, 56 amino acids (14.32%) formed strands, 22 amino acids (5.63%) formed β-turns, and 229 amino acids (58.57%) formed random coils (Figure 3).

**Fig. 3.**
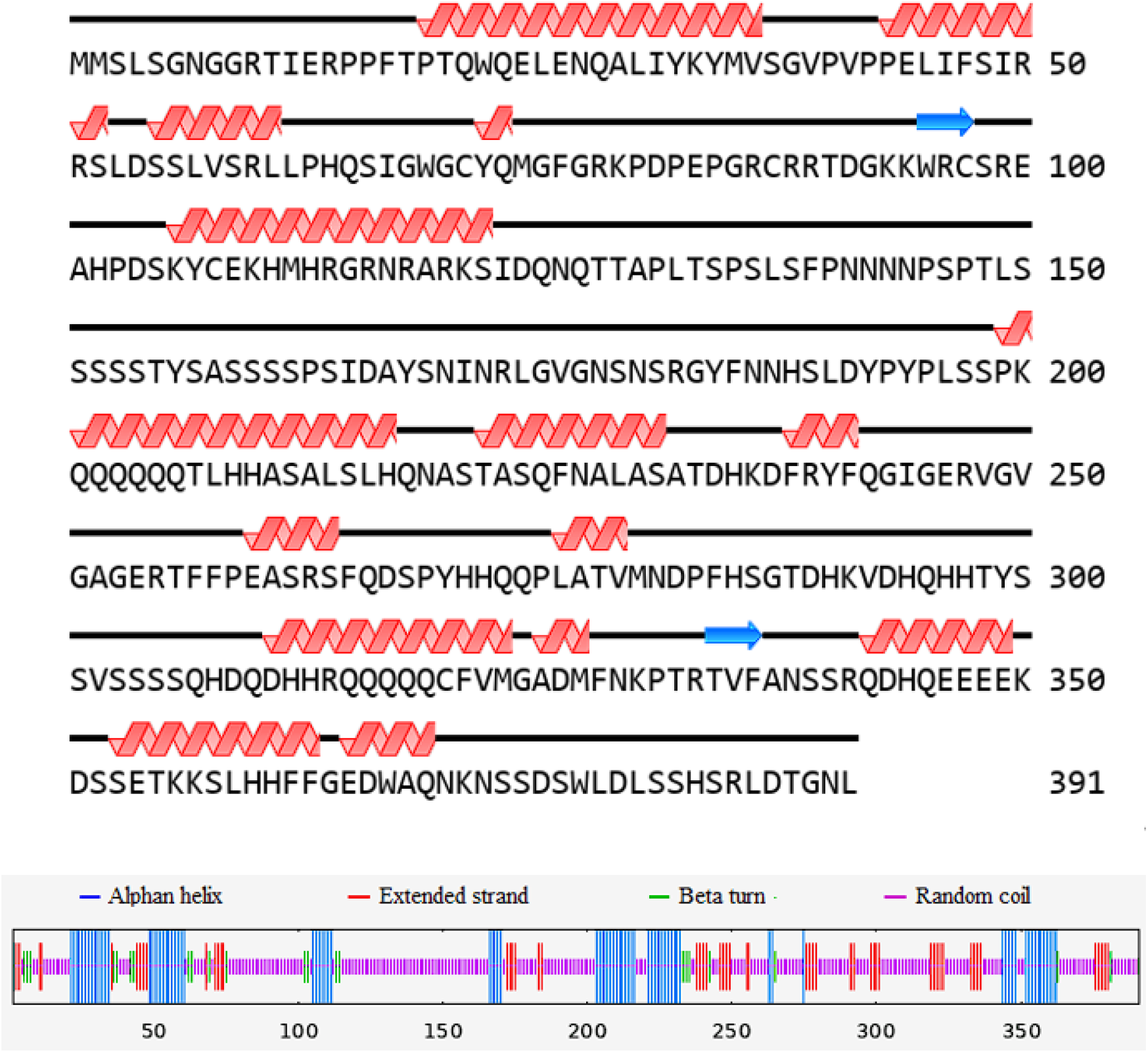
The second structure prediction of BraGRF5. Red spiral indicates Helix and the blue arrow indicates Strand. The blue line indicates Alphan helix, the red line indicates Extended strand, the green line indicates Beta turn and the purple line indicates Random coil.

### GO annotations

The Gene Ontology(GO) database (http://www.geneontology.org/), established by the Gene Ontology Consortium, provides an internationally standardized system for the classification of gene function. Its regulated vocabulary ensures consistency in the description of gene products across various databases by detailing the properties.

GO analysis results revealed that BraGRF5 was particularly enriched in molecular function (MF) such as ATP binding, biological process (BP) such as regulation of transcription, and cell component (CC) such as nucleus (Table 1). These findings suggest that BraGRF5 is involved in complex biological function.

**Table 1.**
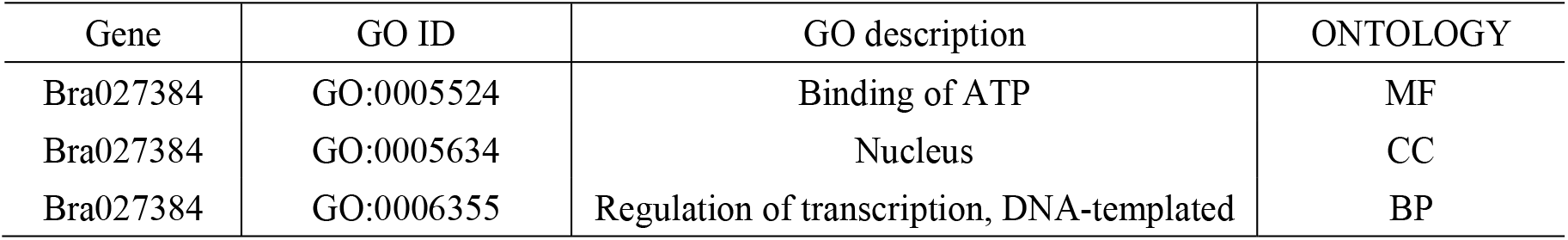
Gene ontology annotation of BraGRF5.

### Prediction of protein-protein interaction networks

The conserved gene network controls the coordinated development of plant organs (Kim & Kende, 2004; Liebsch & Palatnik, 2020). To investigate the potential biological mechanisms of BraGRF5, we predicted its interacting proteins and analyzed their protein-protein interaction networks using STRING database (Figure 4). The BraGRF5-interacting protein had three function domains, auxin response factors (ARFs) required for specific binding to the auxin-response elements (AuxREs) 5’-TGTTCC-3’ for regulating the expression of some ‘early’ genes, C2H2-type zinc finger protein for binding with DNA/RNA/proteins to participate in growth and development as well as the stress response, and ELM2 domain for participating in DNA damage response to repair DNA lesions (Table 2). In light of this, it is speculated that BraGRF5 is involved in multiple biological processes through its interactions within protein-protein interaction networks.

**Fig. 4.**
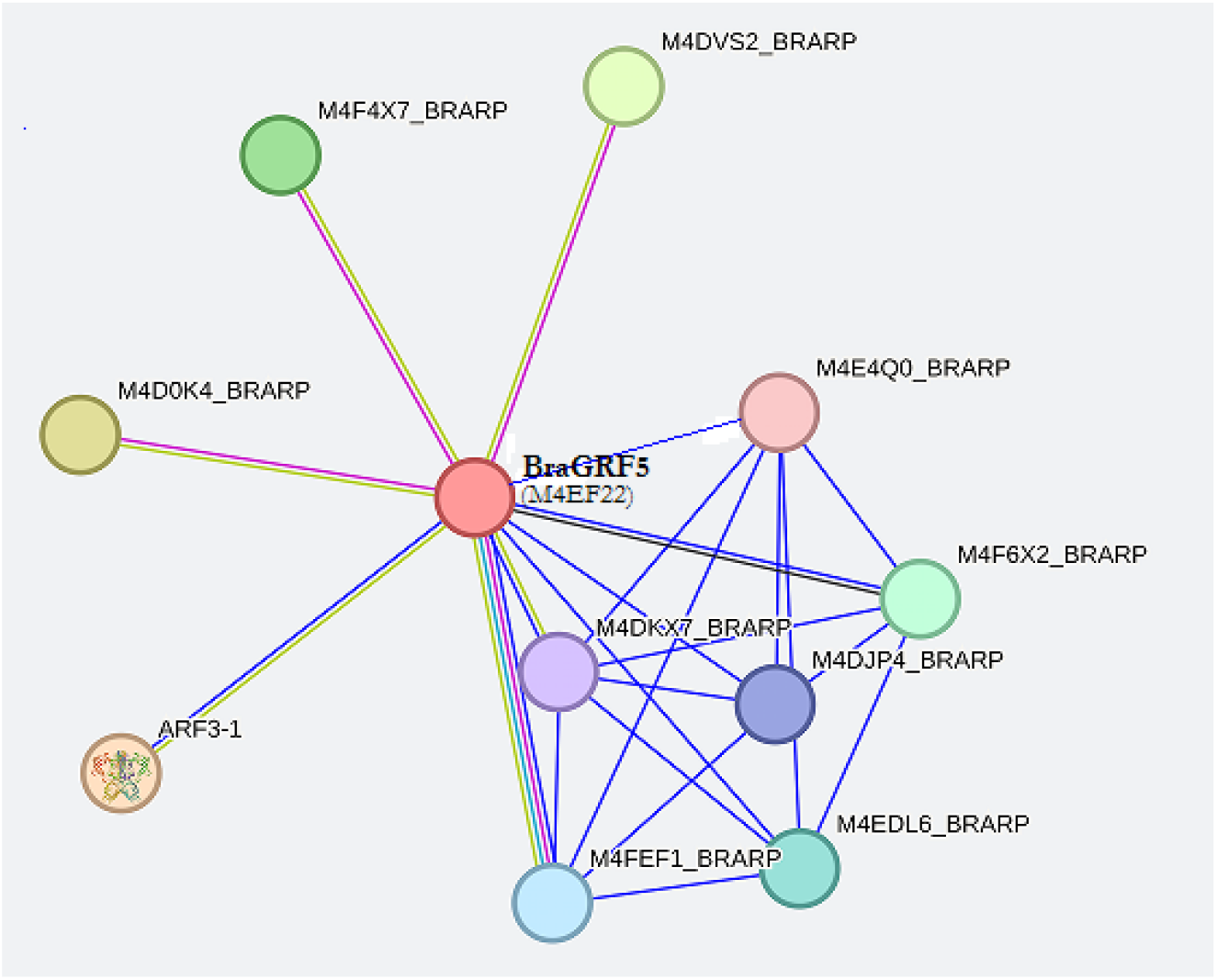
Protein*-*protein interaction networks of BraGRF5 constructed by the STRING.

### Expression profiles of BraGRF5 in different tissues

Based on BRAD, *BraGRF5* exhibited a global expression pattern in *B. campestris*, suggesting a wide range of biological functions. The expression levels of *BraGRF5* varied among tissues, with lower levels leaves, and roots, and the lowest in callus. Conversely, higher expression levels were detected in flowers, siliques, and stems, with the highest expression observed in stems. Notably, *BraGRF5* expression was 314-fold higher in stems compared to callus, and 2.8-fold higher than in siliques (Figure 5).

**Fig. 5.**
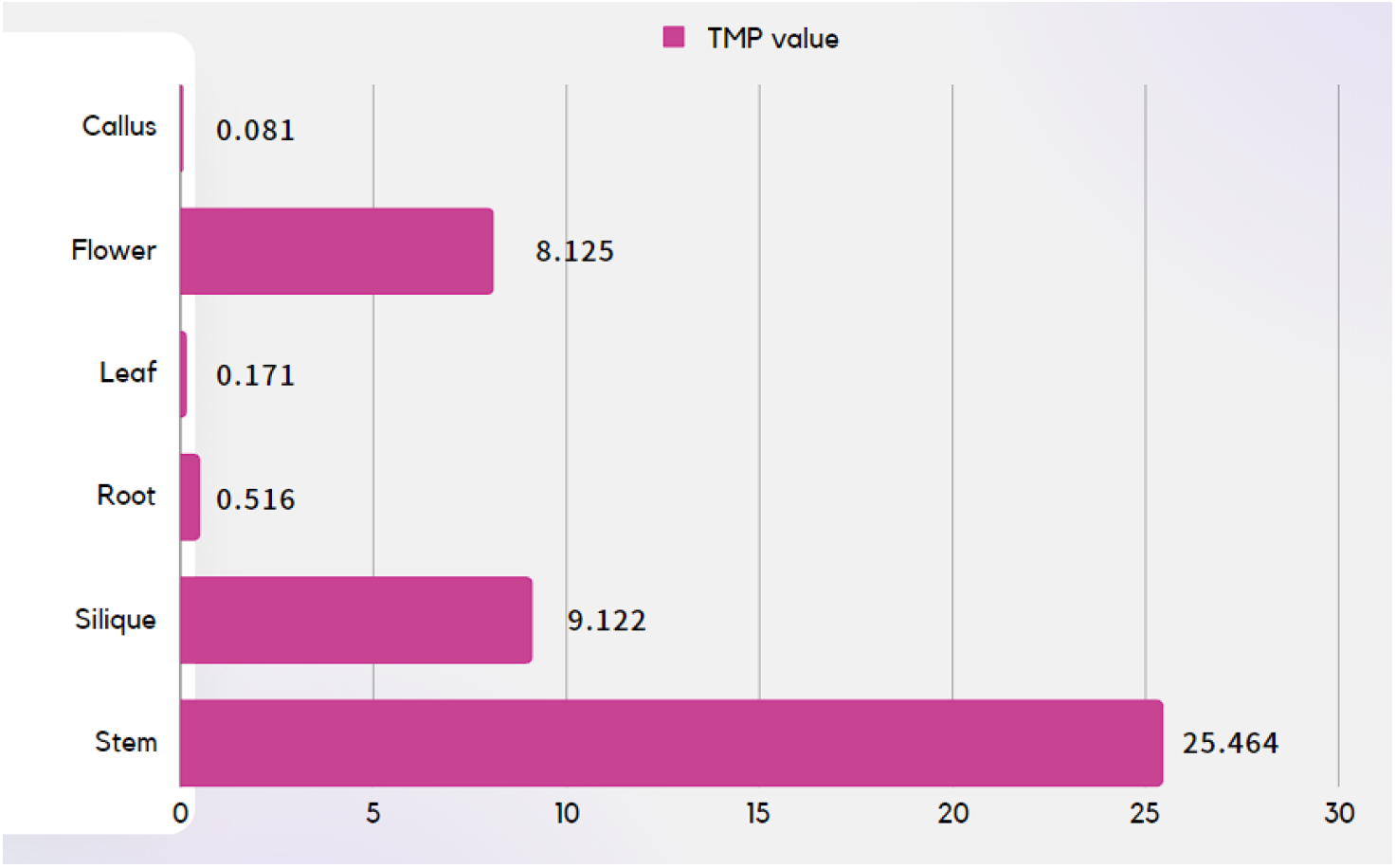
Transcriptome analysis of *BraGRF5* in various tissues. TPM, transcripts per kilobase million

### Chromosomal localization and genome structure of BraGRF5

The *BraGRF5* gene was located on the negative strand of chromosome A05, between 21,359,867 and 21,361,415 bp. This gene encoded 1,549 nt mRNA, containing three exons, 2 introns, and a 1,176 bp coding sequence region (CDS) (Figure 6).

**Fig. 6.**
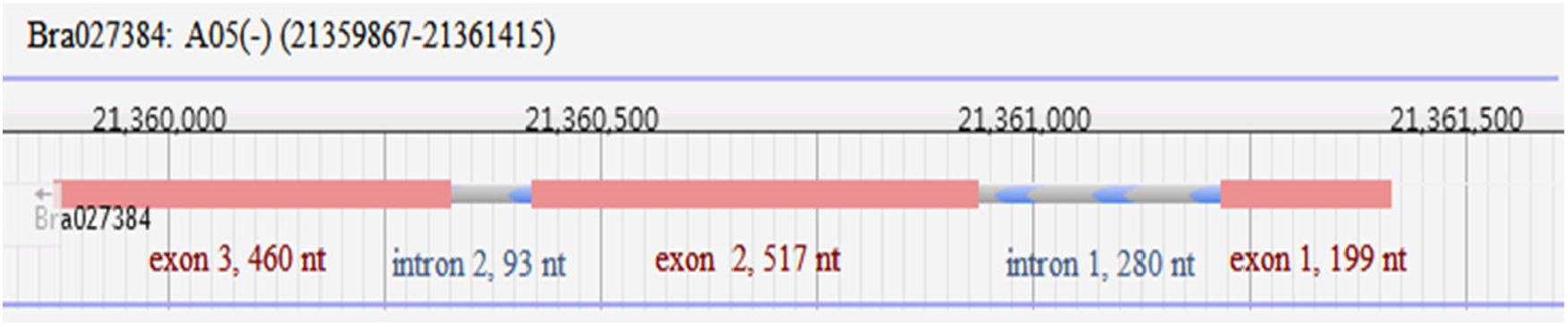
Genomic constructure of *BraGRF5* gene sequence.

### Promoter active region and cis-acting element

Gene expression is regulated by upstream promoters, which play a crucial role in initiating transcription. To explore the regulatory elements of *BraGRF5*, the 2,000 bp region upstream from its start codon was analyzed. Four transcriptionally active promoter regions with Score > 0.8 were predicted by Neural Network Promoter Prediction (Table 3), suggesting their potential promoter function.

**Table 3.**
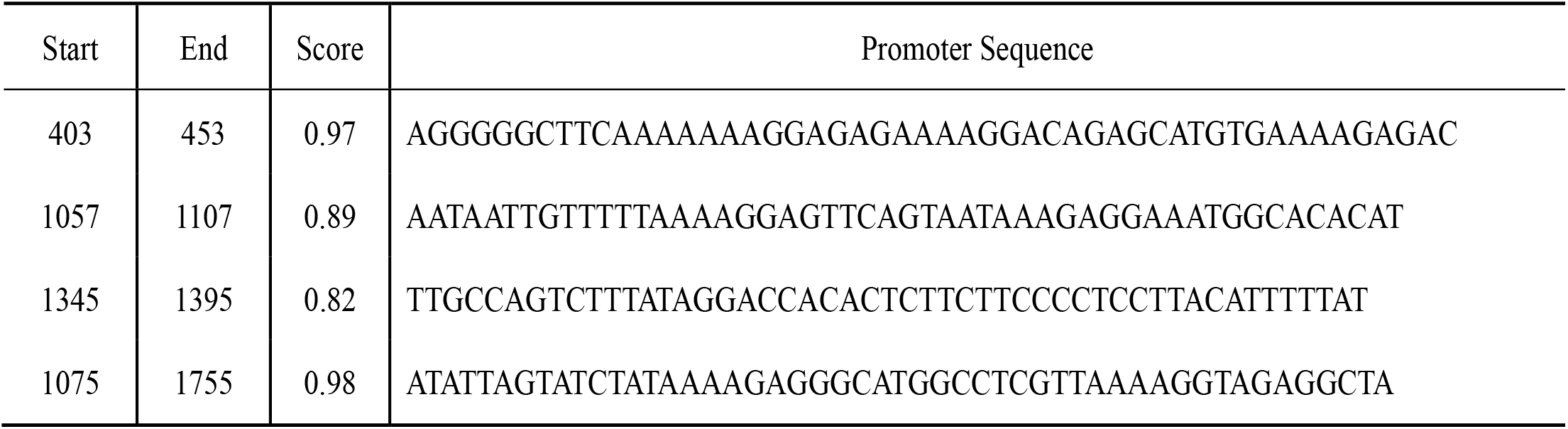
The promoter predictions for *BraGRF5*.

The timing, geographic distribution, and level of gene expression are all significantly influenced by the *cis*-acting regions located within the promoter sequence. Utilizing PlantCare, we found that the *BraGRF5* promoter sequence contained various cis-elements, including light-responsive *cis*-elements such as AE-box, ATC-motif, and GT1-motif. Additionally, hormone-responsive elements for jasmonic acid (JA), gibberellin, salicylic acid, auxin, *etc* were identified, including CGTCA-motif, GARE-motif, P-box, TATC-box, TCA-element, TGA-element, and TGACG-motif. Stress-responsive *cis*-elements, such as the Myb binding site (MBS) involved in drought stress response, as well as TC-rich repeats for defense and stress responsive *cis*-elements, were also present. Furthermore, the promoter sequence contained elements regulated by flavonoid biosynthetic genes, such as MBS, along with canonical core promoter elements like TATA-box and CAAT-box (Table 4). These findings indicate that, in addition to light, *BraGRF5* is regulated by multiple hormones and can respond to various stresses.

**Table 4.**
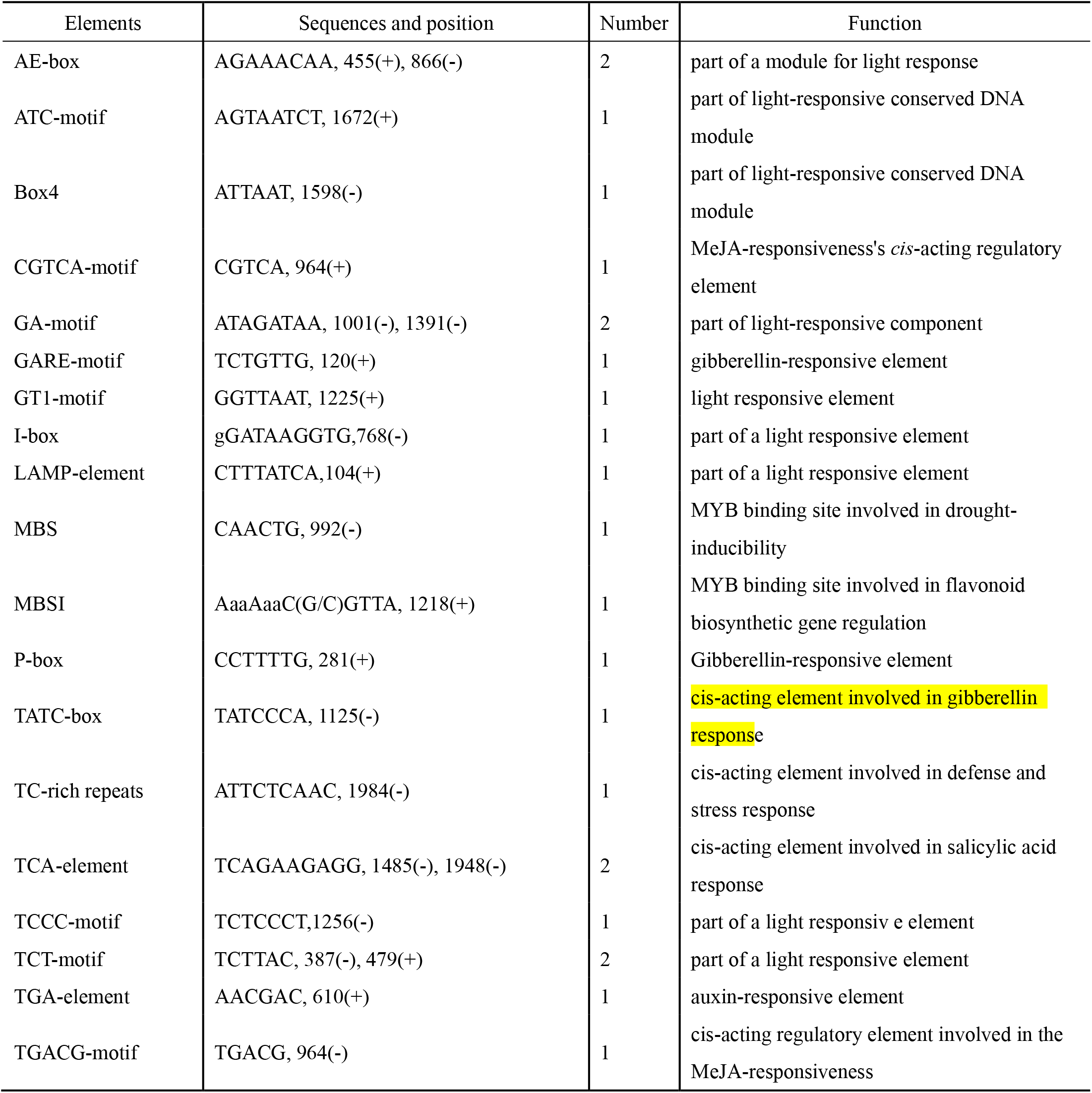
Predicted *cis*-regulatory elements in *BraGRF5* promoter.

## Discussion

The totipotency of plant cells is a remarkable characteristic that enables significant regenerative potential in response to injuries or environmental challenges. These characteristic forms the basis for the establishment of plant asexual propagation and tissue culture techniques (Ikeuchi *et al*., 2016, Mendez-Hernandez *et al*., 2019). In practical applications, the efficiency of tissue culture regeneration-essential for many molecular genetics and breeding techniques-varies significantly across different plant species and genotypes. This variation poses a major limitation to the widespread application of plant regeneration techniques, including plant transformation and genome editing (Altpeter *et al*., 2016).

Genome editing, also known as genetic engineering, is a widely used tool for gene function research and plant breeding, which includes two main stages. The initial stage of genetic modification involves introducing genetic material into plant cells, with two widely used methods being particle bombardment and *Agrobacterium*-mediated transformation; the subsequent stage involves obtaining transformants through tissue culture. However, the efficiency of regeneration and transformation is highly genotype-dependent, which limits the success of genetic engineering, in some crops, gene editing. Developing efficient regeneration and transformation systems remains a significant challenge for enhancing certain plant species through genetic modification.

Recent studies have identified growth regulatory factors (GRFs) as key players in enhancing regeneration and transformation efficiency. Research has shown that a fusion protein combining GRF4 from *T. aestivum* and *GRF-interacting factor 1 (GIF1)* can significantly boost regeneration and genetic transformation efficiency in *T. aestivum, O. sativa*, and other species of *gramineae* family (Debernardi *et al*., 2020).

Growth regulatory factors (GRFs) significantly impact plant growth and development. For instance, overexpression of *OsGRF4* in *O. sativa* increases grain size and panicle length (Sun *et al*., 2016). In *Z. mays*, overexpressing *ZmGRF10* reduces cell proliferation, leading to smaller leaves (Wu *et al*., 2014), whereas in *Tacamahaca* (*P. pseudo-simonii* × *P. nigra* ‘Zheyin 3#’, Ppn), increased expression of *PpnGRF5-1* results in larger leaves (Wu *et al*., 2021). In *A. thaliana, AtGRF1/2* overexpression delays peduncle formation (Kim *et al*., 2003), while *AtGRF5* enhances cell proliferation in leaf primordia, resulting in increased leaf size (Horiguchi *et al*., 2005). Furthermore, transferring seven *BrGRFs* from *B. rap* into *A. thaliana* produced transgenic plants with larger and more abundant cotyledons, leaves, flowers, roots, as well as larger seeds and appendages compared to wild type plants (Hong *et al*., 2017).

Studies have demonstrated that *AtGRF5* from *A. thaliana* promotes cell proliferation in the sprouts of *B. vulgaris*, enhances transformation efficiency, and significantly improves stable transformant production in various crops. Notably, overexpression of endogenous *GRF5-like* genes in crops such as *Helianthus annuus* and *Z. mays* demonstrates even greater effectiveness than the use of exogenous *AtGRF5*. For example, *H. annuus* with overexpressed *HaGRF5-LIKE* exhibit higher GFP expression levels than those with *AtGRF5*, while *Z. mays* overexpressing *ZmGRF5-LIKE* forms embryogenic calli more readily compared to those with *AtGRF5*. These findings suggest that utilizing endogenous *GRF5-like* genes provides better regeneration and transformation potential in these crops (Kong *et al*., 2020).

This study examined the molecular characteristics of *BraGRF5*, the homolog of *A. thaliana AtGRF5* in *B. campestris*. Future research will focus on analyzing the codon bias and other characteristics of *BraGRF5* to provide valuable insights for its potential application in breeding and genetic research. According to the Brassicaceae Database, 17 members of the *GRF* gene family have been identified in *B. campestris*. However, there is no uniform naming convention for these genes. For instance, they are referred to as *BrGRF1–BrGRF17* based on their chromosomal locations and order, while 15 of these members are also named as *BraGRF01– BraGRF15* (Wang *et al*., 2014, Wu *et al*., 2023). This study identified *Bra027384* as the ortholog of *AtGRF5* in *B. campestris* based on sequence similarity, and it has been provisionally designated as *BraGRF5*. However, the naming convention for the *BraGRF* gene family in *B. campestris* still requires further standardization.

Gene expression is mainly regulated by the promoter sequence located upstream of the gene. While the promoter does not encode proteins, it controls gene activity through interactions between *cis*-acting elements and trans-acting factors. Gibberellic acid (GA) has been shown to induce the expression of *OsGRF1*, promoting stem elongation of *O. sativa* (van der Knaap *et al*., 2000), to increase the expression of *DaGRF2* in *Dioscorea alata* (Xing *et al*., 2020), and to boost the expression of nine *BrGRFs* by more than fivefold (Wang *et al*., 2014). Additionally, GA and jasmonic acid (JA) upregulate the expression of *SlGRF2* and *SlGRF5* in *S. lycopersicum*, respectively (Khatun *et al*., 2017).

GRFs respond dynamically to various stresses. For example, dark stress reduces *GmGRF* expression in leaves of *G. max* (Chen *et al*., 2019), while in *A. thaliana, AtGRF5* interacts with cytokinin to promote cell division (Vercruyssen *et al*., 2015). In drought-stressed *S. lycopersicum* plants, *SlGRF13* expression level decreases, while *SlGRF6* expression level increases, indicating that different *SlGRFs* play distinct roles in response to drought (Khatun et al., 2017).

In this study, we identified a 2,000 bp upstream region of the *BraGRF5* transcription start site in *B. campestris*, which exhibited transcriptional activity and contained multiple regulatory elements, including light- and hormone-responsive cis-elements. These included elements responsive to JA, gibberellin, salicylic acid, and auxin. This suggests that modifying environmental factors, such as light exposure and hormone levels, could enhance *BraGRF5* expression, potentially improving tissue culture outcomes and transformation efficiency. Precise control of *BraGRF5* expression through environmental factors may provide an alternative to transgenic approaches, circumventing challenges such as random chromosomal integration and genotype dependency that are often associated with genetically modified crops.

## Conclusion

The analysis of *BraGRF5* offers valuable insights into its potential applications in molecular breeding and genetic transformation of *B. campestris*. Further research into its molecular characteristics, the standardization of naming conventions for the *BraGRFs* family, and an understanding of its regulatory mechanisms under various growth conditions will advance molecular breeding and transformation technologies in *B. campestris*. These findings lay a foundation for enhancing crop development and transformation efficiency.

## FUNDING

The Basic Research Program of Shanxi Province (20210302124141) and the Applied Basic Research Program Project of Shanxi Province, China (202103021224148) provided funding for this work.

## Notes

### Competing Interest Statement

The authors have declared no competing interest.

